# HIGH-FIDELITY BACKPROPAGATION THROUGH PRIMATE FOVEAL CONES

**DOI:** 10.64898/2026.01.28.701353

**Authors:** Sophia R. Wienbar, Gregory S. Bryman, Michael Tri H. Do

**Author notes:** Correspondence: Sophia R. Wienbar and Michael Tri H. Do. Equal contribution (Sophia R. Wienbar, Gregory S. Bryman).

## Abstract

Primate vision has exceptionally high spatial acuity and contrast sensitivity. This performance originates in specialized photoreceptors of the fovea. These cones transduce light into electrical signals in the outer segment, and convey these signals to the presynaptic terminal for transmission. Backpropagating signals are also possible, as the terminal receives inputs. Such signals could influence phototransduction itself. To test this idea, we recorded electrophysiologically from both ends of single cones dissociated from the macaque fovea. We found that backpropagation was effective despite the extreme slenderness and length of these cells. Backpropagation was also effective in a passive compartmental model, indicating that amplification by voltage-gated channels is not required. We then modeled foveal cones receiving terminal inputs from retinal networks. Despite faithful backpropagation of these inputs, they appear unlikely to influence phototransduction. Thus, even though foveal cones exhibit effective backpropagation, their encoding of visual information may remain compartmentalized.

**SIGNIFICANCE STATEMENT:** Humans, like other primates, see a fineness of detail that eludes other mammals. This capability is used for tasks like reading and recognizing faces. It is lost in leading forms of vision impairment, such as age-related macular degeneration. Investigating it therefore provides insight into the origin of exceptional sensory performance while strengthening the foundation for preserving and restoring sight. This study examines cells that initiate high-acuity vision, the foveal cones, which produce electrical signals from light and send them forward. It reveals that electrical signals also travel effectively in reverse, from the site of transmission to that of production, and how the production of light responses may be independent nonetheless.

## INTRODUCTION

The classic neuron processes signals, sends them forward through its axon, and transmits them to other cells through its presynaptic terminal. Many neurons also receive inputs at their presynaptic terminals, which may trigger signals that backpropagate through the axon to influence signal processing (Stuart et al., 1997; Waters et al., 2005; Alle and Geiger, 2008; Paradiso and Wu, 2009; Debanne et al., 2011). Backpropagation has been studied largely in cortex, where it mediates important processes that include setting synaptic strength (Waters et al., 2005). Less is known about backpropagation in primary sensory neurons, where it has the potential to shape the environmental information that is available to the organism.

Conscious vision in humans and other primates relies on photoreceptors of the fovea, a specialization of the central retina (Provis et al., 2013). These foveal cones are uniquely narrow and densely packed, forming an array that resolves fine spatial details in the image. Primates see a greater level of detail than other mammals, which lack foveas; for example, ten-fold higher than cats and a hundred-fold higher than mice (Caves et al., 2018). A foveal cone converts light into electrical activity in the outer segment (OS), and conveys it through a long axon (up to ∼400 µm) to the presynaptic terminal (Perry and Cowey, 1988; Bryman et al., 2020). Cone terminals are understood to receive signals, including gap-junctional currents from neighboring cones and negative feedback from horizontal cells (Boycott and Kolb, 1973; Ahnelt and Pflug, 1986; Tsukamoto et al., 1992; DeVries et al., 2002; Verweij et al., 2003; Hornstein et al., 2004; O’Brien et al., 2012; Kim et al., 2025). In the OS, Ca^2+^ flux through the ion channels of phototransduction sets the adaptation state of the cell (Yau, 1991; Burns and Baylor, 2001). If backpropagated terminal signals changed the driving force for Ca^2+^ enough, they could influence the initiation of high-acuity vision.

Foveal cones signal using graded voltages rather than regenerative action potentials (Sinha et al., 2017; Bryman et al., 2020; Saha et al., 2024). Forward propagation of these analog signals is highly effective (Bryman et al., 2020). Even though foveal cones are thin and long, their intracellular resistivity is low and their membrane resistivity is high, which allows current to flow freely within the cell with little loss to the extracellular space. These passive properties appear to be sufficient and nearly optimal for propagation. Indeed, within the natural range of foveal cone lengths, most signal frequencies propagate forward with <10% attenuation. These observations indicate that analog backpropagation through foveal cones should be effective, though the idea is untested.

Much is unknown about the physiology of foveal cones. To a large degree, this is because of availability; the fovea is just a percent of the primate retina (Provis et al., 2013). This small size belies the fovea’s importance. It is represented by up to a quarter of primary visual cortex (Born et al., 2015). A few millimeters outside the fovea, humans are legally blind (Masland, 2017). Age-related macular degeneration, which selectively affects the central retina, is a leading form of vision loss (Fleckenstein et al., 2024). Investigations of the fovea provide basic knowledge—for example, concerning the basis of high visual acuity and the influence of a widespread process like backpropagation—and may support advances in treating blindness. Here, we investigate the directionality of information flow in foveal photoreceptors and its potential influence on encoding light.

## MATERIALS AND METHODS

### Materials Availability

This study did not generate new unique reagents.

### Data and Code Availability

NEURON models and analysis routines are available at github.com/wienbar/ConeBackpropagationCode_2026. Raw data are available upon request.

### Experimental model and subject details

All procedures were approved by the Animal Care and Use Committees of institutions that provided tissue and Boston Children’s Hospital. Macaques (*Macaca mulatta*, 3-14 years of age; *Macaca fascicularis*, 3-9 years of age) of both sexes were used. Animals were euthanized for purposes unrelated to the present study. Most had prior experimental histories, but none had known vision problems. No findings reported here were observed to covary with animal age, sex, or experimental history.

### Tissue collection

Eyes were removed pre-mortem under deep anesthesia. Post-mortem eyes were also used, in which case the ischemic time was usually <15 min but never >60 min. The eye was hemisected coronally, vitreous humor removed mechanically, and the posterior eyecup immersed in oxygenated media (see below). These procedures were completed within 5 minutes. Eyecups were transported in darkness to the laboratory (15-90 min). Experimental observations appear independent of tissue ischemic time and time elapsed since eye removal.

### Solutions

External solutions had osmolarities of 280-285 mOsm unless otherwise noted. Bicarbonate-buffered solutions were supplemented with penicillin (80-96 U/ml) as well as streptomycin (0.080-0.096 mg/ml; Millipore-Sigma P4333) and equilibrated with carbogen (95% O_2_ and 5% CO_2_) for a pH of 7.4. The standard external solution was bicarbonate-buffered Ames medium (“bicarbonate Ames”; Millipore-Sigma A1420). For tissue transport and culture, the glucose concentration of Ames was sometimes increased from 6 to 24 mM. For microdissection, tissue perfusion sometimes used a reduced, “ionic Ames” medium (Do et al., 2009): in mM: 120 NaCl, 3.1 KCl, 0.5 KH_2_PO_4_, 1.2 CaCl_2_, 1.2 MgSO_4_, 6 glucose, and 22.6 NaHCO_3_. “HEPES Ames” was ionic Ames buffered with 10 mM HEPES rather than bicarbonate (with NaCl increased to 140 mM; pH 7.4 with NaOH). Trituration solution (Do et al., 2009) contained (in mM) 70 Na_2_SO_4_, 2 K_2_SO_4_, 10 glucose, 85 sucrose, 5 MgCl_2_, and 10 HEPES (pH 7.4 with NaOH; 305 mOsm). The pipette (electrode) solution contained (in mM) 97 K-methanesulfonate, 13 NaCl, 2 MgCl_2_, 1 CaCl_2_, 10 EGTA, 10 HEPES, 0.3 Na-GTP, 4 Mg-ATP, 7 phosphocreatine di(tris), and 2 L-glutathione (pH 7.2 with KOH, bringing [K^+^] to 118 mM; 275 mOsm).

### Tissue culture and microdissection

All procedures on live tissue were performed under infrared illumination. To remove any residual vitreous humor, human plasmin was sometimes applied (Stalmans et al., 2010)(Sigma P1867; 1-2 µM in Ames medium, 23 or 35 °C, 20-30 min). Eyecups were dark-adapted for >1 hr at 33 °C before experiments started and were kept for ≤48 hrs total. Tissue was removed from the eyecup as needed.

### Isolation of cone photoreceptors

Cones were dissociated from the retina using procedures described previously (Bryman et al., 2020). The fovea (centered in a ∼5-mm^2^ piece of retina) or peripheral retina (≥8 mm from the fovea, also taken in a ∼5-mm^2^ piece) was dissected from the sclera and sometimes retinal pigment epithelium. Tissue was incubated for 15 min in papain (Worthington LS003119, 20-31 U/ml; L-cysteine from Millipore-Sigma W326305 was usually omitted but sometimes included at 3 mM; 35 °C), then washed in Ames medium (23 °C). This medium often contained 1 mg/ml ovomucoid trypsin inhibitor and 1 mg/ml bovine serum albumin (Worthington LS003087 and Millipore-Sigma A8806, respectively). In some cases, DNase I was included in the incubation or washes to reduce cell clumping (Millipore-Sigma DN25 at 120-200 U/ml or Worthington LS006333 at 120 U/ml). Tissue was transferred into trituration solution and passed through a series of fire-polished glass pipettes that had decreasing bore diameter. An aliquot of the cell suspension was placed in the recording chamber and mixed with HEPES Ames. After cells adhered, bicarbonate Ames was superfused (∼2 ml/min).

### Electrophysiological recordings

Cells were visualized with a 60× water-immersion objective (1.0 NA), differential interference contrast optics, and infrared transillumination (center wavelength of 850 or 940 nm). Whole-cell recordings were made at room temperature (23 °C) for increased stability. They employed a Multiclamp 700B amplifier with a 4-pole, low-pass Bessel filter (10 kHz). The sampling rate exceeded the Nyquist minimum. Pipettes were borosilicate glass (A-M Systems 603500) and wrapped with parafilm to reduce capacitance. Seals (generally ≥10 GΩ) were formed on the IS and terminal. Whenever possible, seal stability was confirmed by pulling an outside-out patch after recording. Pipette resistances (3-12 MΩ) were matched between IS and terminal for individual cells. Series resistance (R_s_) was typically <50 MΩ. A 6.5-mV junction potential has been corrected (Neher, 1992). For experiments involving current injection, the voltage drop across R_s_ was corrected offline. Recordings were terminated or excluded from analysis if changes in cellular morphology appeared; if R_s_ was ≥50 MΩ or varied by ≥50%; or if the voltage drift (generally <3 mV) was >10 mV. No dependencies of data on R_s_ were observed.

### White-noise electrical stimulation and filter construction

Gaussian white noise stimuli (sampled at 10 kHz) were set to drive each cell across the physiological range of photoreceptor light responses (roughly −40 mV to −70 mV in the various species examined)(Barnes, 1994; Okawa et al., 2008). The currents used had mean offsets as large as −50 pA and standard deviations as large as 450 pA. Stimuli typically lasted 50 s. The first 10 s allowed any response transients to settle and were not analyzed. The next 30 s were used to compute the linear filter as the cross-correlation of the stimulus and the response, normalized by the power spectrum of the stimulus (Kim and Rieke, 2001; Baccus and Meister, 2002). The cross-correlation was computed in 5-s windows with a 0.5-s overlap and averaged. The last 10-s period was not used for filter construction. It was convolved with the filter to give a predicted response for comparison with the actual, measured response. The filters were examined in the frequency domain to quantify response magnitude and phase lag (relative to the stimulus) as a function of temporal frequency. Changes in the offset and standard deviation of injected currents had only a mild effect on filter shape (i.e., small changes limited to <20 Hz) and no detectable effects on filter linearity and propagation fidelity.

### Compartmental modeling

Passive, compartmental models were generated in NEURON using its standard integration method (backward Euler integration)(Carnevale and Hines, 2009). Cells had 4-5 sections (IS, soma, axon, terminal, and sometimes an OS). Each section was subdivided into segments whose number followed the conventional rule of at least 10 per approximated alternating current length constant (λ_AC_). This is an approximation because it assumes that the resistive term is negligible (Carnevale and Hines, 2009). The frequency was set to a conservative value (≥500 Hz). Increasing the number of segments had no apparent effect on responses. Time steps were generally 10 µs. Membrane conductance was passive and had a linear (i.e., ohmic) current-voltage relation. The extracellular space was non-resistive and non-capacitive.

### Reference foveal cone model

The “reference” foveal cone had compartments of (length×diameter in µm): IS, 30×3.5; soma, 5.5×5.5; axon, 400×1.6; and terminal, 7×4.5. These parameters are characteristic of the longest and most slender foveal cones, observed within a few hundred microns of the center of the fovea (Perry and Cowey, 1988; Curcio et al., 1990; Hsu et al., 1998; Drasdo et al., 2007). The specific membrane capacitance was initialized to 1 µF/cm^2^, the axial resistivity to 76.8 Ω·cm, and the membrane resistivity to 1.2×10^4^ Ω/cm^2^. All of these parameters were derived from morphological and electrophysiological measurements (Bryman et al., 2020).

### Outer segment model

Outer segments were modeled using existing morphological measurements (Dowling, 1965). The outer segment had a length of 40 µm and a diameter of 0.9 µm, with two sealed ends because the connecting cilium has a minuscule diameter. Each disc had a thickness of 0.1 µm and a diameter of 0.9 µm. Each outer segment contained 1,200 discs (Dowling, 1965). Cohen estimated that 10% of discs are continuous with the plasma membrane (Cohen, 1961; Dowling, 1965). Because points of continuity could have been missed in these analyses of single image planes, 60% continuity was also tested. The surface area of continuous discs was added as a lengthening or broadening of the outer segment. The modeled outer segment had the same membrane capacitance, intracellular resistivity, and membrane resistivity as the other compartments.

### Gap-junctional network model

Networks of cones were formed by connecting the cone terminals in a hexagonal array with gap junctions. The gap-junction conductance implementation was as described in the NEURON documentation (Carnevale and Hines, 2009). An array of 37 cones was used. Other array sizes were tested (7-61 cones) and no qualitative differences found. Voltage spread is minimal (<8%) beyond the first neighboring cone for coupling conductances of <2 nS; this and larger conductances are well beyond the estimated physiological range (DeVries et al., 2002; Hornstein et al., 2004), though an upper bound of 10 nS was suggested by an early study (Tsukamoto et al., 1992).

### Center and surround model

This simulation used light-evoked currents recorded from macaque peripheral cones by Schnapf and colleagues (Verweij et al., 2003). These currents were measured for illumination of the center (a spot with a diameter of 35 µm) or surround (an annulus with an inner diameter of 40 µm and an outer diameter that varied from 75 to 350 µm). Center currents were fit with a sum of five Gaussian functions:

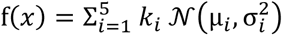

Surround currents were fit using a flat-top Gaussian function:

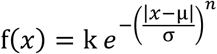

These fits were used in simulations with the reference foveal cone model. The center current was played into the outer segment and the surround currents were played into the terminal. The center and surround currents were aligned in time by the onset of illumination.

### Experimental design and statistical analyses

Because effect sizes and variances had yet to be defined, experiments were designed, performed, and analyzed iteratively. The final dataset includes 23 cells from 12 animals, with 1-4 cells collected per animal. Data were analyzed in Python (v3.11.19). NEURON (v8.2.2) modeling was also performed in Python. Figures were assembled in Adobe Illustrator (v27.5). For statistical tests, the Wilcoxon signed rank test was used to compare nonparametric, paired data. Except for baseline current and input resistance (**Figure 1C, F** and **Tables S1** and **S2**), three tests were performed per parameter measured and the Bonferroni p-value correction factor was 3.

**Figure 1.**
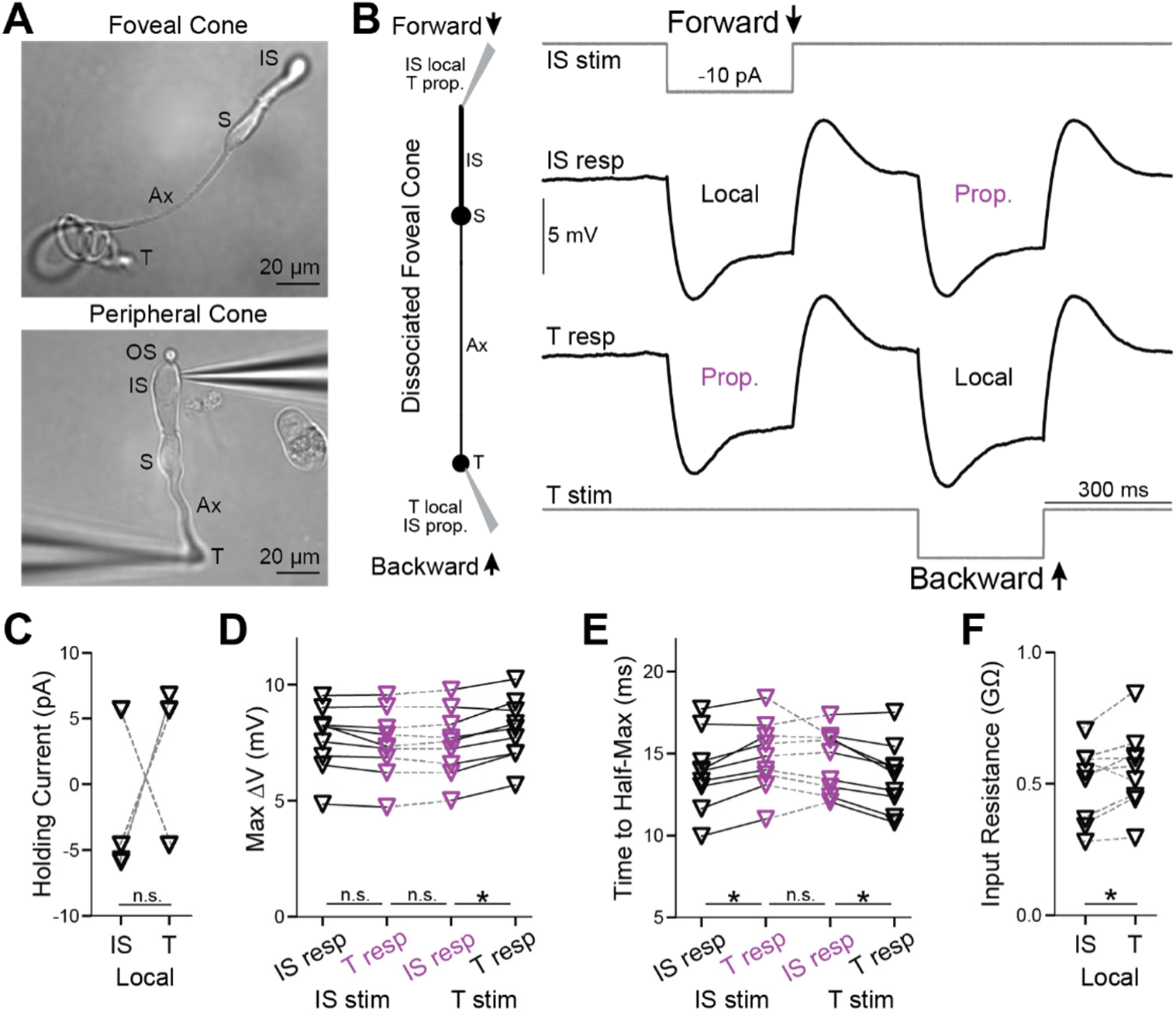
Backpropagation of responses to steps in foveal cones. **A.** Live, acutely dissociated foveal (top) and peripheral (bottom) cones, the latter with electrodes attached. Labeled are the outer segment (OS), inner segment (IS), soma (S), axon (Ax), and terminal (T). **B.** *Left,* Schematic of a dissociated foveal cone, its compartments (inner segment, IS; soma, S; axon, Ax; and terminal, T), and response nomenclature. *Right,* Simultaneous, whole-cell recordings from a dissociated foveal cone’s IS and terminal. Square pulses of current were injected at the IS to examine forward propagation (top stimulus monitor and adjacent trace) or at the terminal to examine backpropagation (bottom stimulus monitor and adjacent trace). Local (at the injection site) and propagated (distal to the injection) responses are labeled. Traces are averages of 10 trials. Steady currents were injected at the IS and terminal to keep their resting voltages at -60 mV. **C.** Steady currents injected into the IS and terminal of each foveal cone to maintain the membrane voltage at -60 mV. Several symbols overlap, and “n.s.” indicates p≥0.05. **D.** Maximum voltage responses for forward (IS stim) and backward (T stim) propagated responses in foveal cones (asterisk indicates p<0.05 with a Bonferroni correction for 3 comparisons). **E.** As in **D** but for the time taken to reach the half-maximal response. **F.** Input resistances for the IS and terminal of each foveal cone (calculated from local responses). 9 cells; asterisk indicates p<0.05.

## RESULTS

To measure response propagation in macaque foveal cones, we performed electrophysiological recordings from both the inner segment (IS) and axon terminal of individual cells. Doing so in the intact retina is impractical because these structures are tiny, connected pairs must be found within dense arrays of near-identical structures across distances of up to ∼400 µm, and terminals lie beneath tough tissues. Therefore, we acutely dissociated foveal cones from the macaque retina (**Figure 1A**)(Bryman et al., 2020). Dissociation removes accessory tissues and, usually, the delicate outer segments (OS); however, forward propagation appears to be little affected (Bryman et al., 2020). For individual cones, we made simultaneous whole-cell, patch clamp recordings from the IS and terminal. We injected current into the IS or terminal, which ordinarily receives current directly from phototransduction or retinal circuitry, respectively, and measured the voltage at both locations. We compared the voltage of the stimulated compartment (local response) with that of the opposite compartment (propagated response) to assess bidirectional signal flow through each tested cone.

We dissociated peripheral cones for comparison (**Figure 1A**). For all experiment types discussed below, we observed no differences in peripheral cones between forward- and backward-propagating responses, whether considering magnitude or kinetics (**Figure S1**, **Tables S1**-**S4**). This is consistent with the short and broad shape of these cells (Bryman et al., 2020). The remainder of this **Results** section concerns foveal cones.

### Backpropagation of transient and steady-state responses

To facilitate comparison between the IS and terminal of each foveal cone, we maintained both compartments at the same voltage (near -60 mV) by injecting steady current (**Figure 1B**). These currents did not differ systematically between the compartments (9 cells, p=0.82; Wilcoxon signed rank test for paired data unless otherwise noted; **Figure 1C**). We then delivered a step stimulus whose polarity and size resembled that of a modest phototransduction current, and whose duration allowed examination of transient and steady-state voltage changes (-10 pA, 300 ms). This current step evoked a hyperpolarization that was initially large and then relaxed to a plateau; after the step, the response overshot baseline and then settled. These dynamics are expected from vertebrate rods and cones (Barnes, 1994; Van Hook et al., 2019). We found that the response had a similar shape whether local or propagated, and whether initiated at the IS or terminal.

Delivering the current step to the IS, we observed responses whose peak amplitudes at the IS (local) and terminal (propagated) were indistinguishable (7.7 ± 1.4 and 7.4 ± 1.5 mV, 9 cells, p=0.059; when multiple comparisons are made, p values are Bonferroni-corrected; **Figure 1D**, **Tables S1** and **S2**). The rise time to half-maximum (t_1/2_) was mildly faster for the local response than for the propagated response (13.9 ± 2.4 ms and 14.8 ± 2.2 ms, p=0.023, **Figure 1E**). Thus, forward propagation appears to be highly effective when examined with this step stimulus.

For terminal stimulation, the peak amplitudes of local and backpropagated responses differed slightly (8.0 ± 1.4 mV at the terminal and 7.5 ± 1.5 mV at the IS, p=0.023; **Figure 1D**). The t_1/2_ values were 13.6 ± 2.1 at the terminal and 14.6 ± 1.9 ms at the IS (p=0.023; **Figure 1E**). These experiments indicate that backpropagation through foveal cones is effective (see **Tables S1** and **S2** for additional analysis of these and the following experiments).

The amplitude of the propagated response is a smaller fraction of the local response for backpropagation (0.93) compared to forward propagation (0.97). This suggests slightly more loss during propagation from terminal to IS than vice versa. However, propagated responses have indistinguishable amplitudes whether initiated at the terminal or IS (p=1.0). Backpropagation shows a smaller fractional response because the local response has a higher amplitude, consistent with the terminal having a higher input resistance (0.55 ± 0.15 GΩ, compared to 0.50 ± 0.14 GΩ at the IS, p=0.039; **Figure 1F**). Propagated responses also have similar t_1/2_ at the IS and terminal (p=1.0). The amplitudes of the steady-state, propagated responses are also similar (p=1.0). Therefore, when considering the propagated responses themselves, rather than comparing them to local responses, backward and forward propagation appear equally effective.

### Backpropagation of responses to temporal white noise

To examine response propagation across a broad range of temporal frequencies, we injected steady currents to maintain both the IS and terminal near -60 mV and then, at the stimulus site, varied the current about its mean in the form of white noise (**Figure 2A, B**)(Bryman et al., 2020). We measured the resulting voltages at the IS and terminal, then cross-correlated each with the stimulus to generate linear filters (**Figure 2C, D**). These filters provide concise descriptions of the cone responses. Convolving them with the stimulus gave predicted responses that matched measured responses, with no nonlinearity required (Pearson’s R=0.923 ± 0.093, 10 cells; **Figure S2**). Taking magnitude and phase spectra of the linear filters reveals response amplitude and timing as a function of temporal frequency (**Figure 2C, D** and **Figure S3A**)(Kim and Rieke, 2001; Baccus and Meister, 2002; Bryman et al., 2020). We examined absolute magnitude and phase for both forward- and backward-propagating responses. To summarize, for both propagation directions, we found some statistical differences but small effects (**Figure 2E, F**; **Tables S3** and **S4**). For example, propagated responses were indistinguishable at the IS and terminal; assessed at 1, 60, and 100 Hz, p values were 1.0 for magnitude and 0.32-1.0 for phase. Therefore, across temporal frequencies, forward and backward propagation through foveal cones appears to be highly effective.

**Figure 2.**
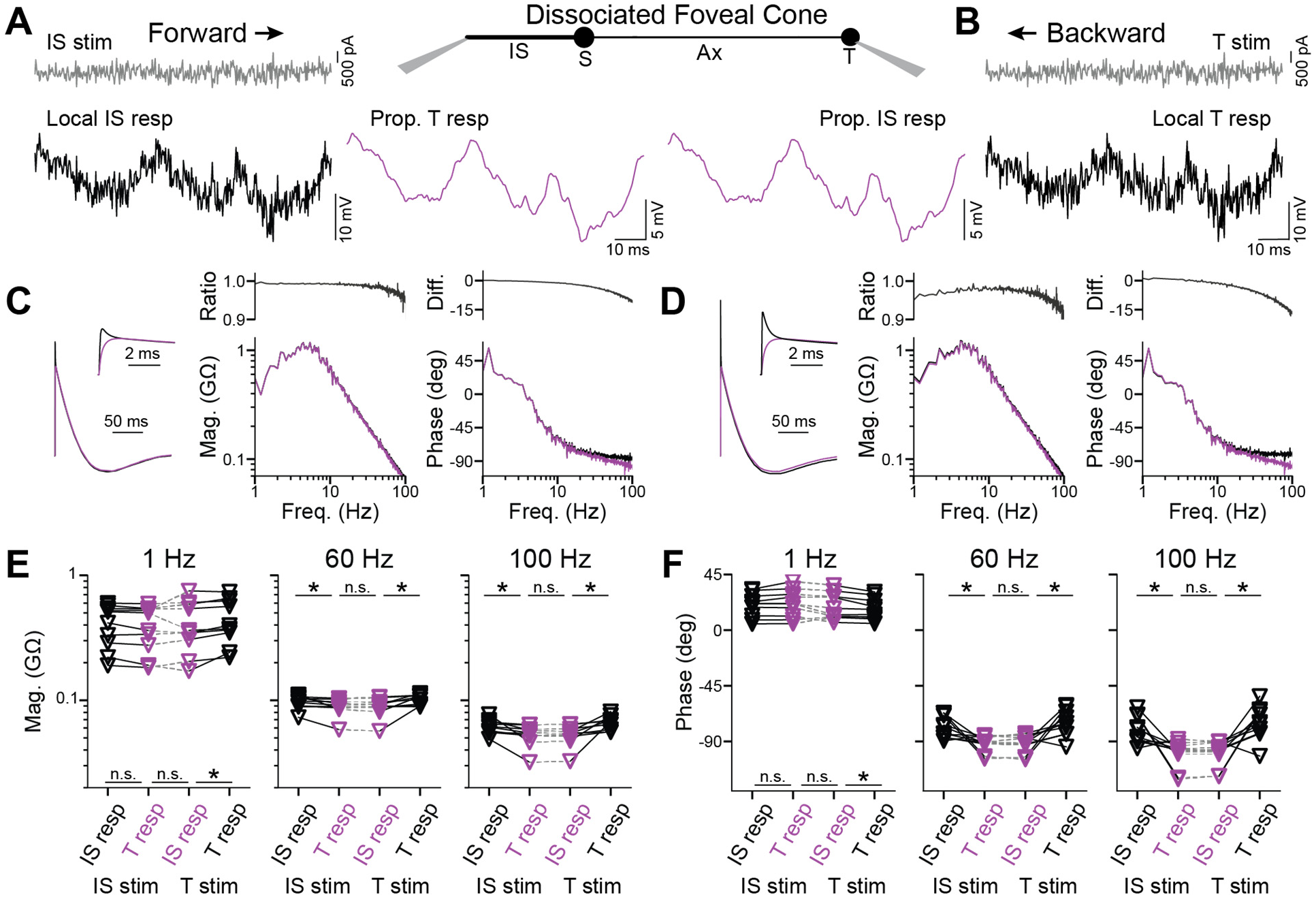
Backpropagation of responses to white noise in foveal cones. **A.** *Center,* Schematic of a dissociated foveal cone and its compartments. *Left,* Current in the form of white noise (gray) was injected into the IS and responses recorded in the IS (black, local) and terminal (magenta, propagated) to examine forward propagation. **B.** Recordings from the same cell as in **A** but for backpropagation, where the current was injected into the terminal and responses recorded in the terminal (black, local) and IS (magenta, propagated). **C.** Linear filters of the IS (black) and terminal (magenta) for forward propagation, alongside their magnitude and phase spectra. The magnitude ratio and phase difference between local and propagated filters are also shown. These filters are calculated for the responses illustrated in **A**. See **Figure S3A** for higher frequencies. **D.** As in **C** but for backpropagation. These filters are calculated for the responses illustrated in **B**. **E.** Response magnitude at 1 (left), 60 (center), and 100 (right) Hz in the IS and terminal for stimulation at the IS (forward propagation) or terminal (backpropagation; 10 cells, asterisk indicates p<0.05 with Bonferroni correction for 3 comparisons). **F.** As in **E** but for response phase.

### Backpropagation in a passive model of a foveal cone

Forward propagation through foveal cones does not require amplification from voltage-gated ion channels and, indeed, is exhibited by a passive compartmental model of the cell (Bryman et al., 2020). We used modeling to determine if passive properties also suffice for effective backpropagation. We used empirical estimates of the specific membrane capacitance, intracellular resistivity, and membrane resistivity of foveal cones (Bryman et al., 2020). To provide a conservative test case for propagation fidelity, we adopted anatomical parameters from the longest and most slender foveal cone that we and others have observed (**Materials and Methods**). We call this model the reference foveal cone. We began by simulating the current step that we used experimentally (10-pA amplitude, 300-ms duration; see above). The voltage reached a plateau during the step and then decayed to baseline afterward (**Figure 3A**). There were no transients, consistent with the absence of active membrane properties (**Figure 1B**). Forward propagation was highly effective in this model (**Figure 3A**), as expected (Bryman et al., 2020). Backpropagation was also highly effective. The propagated responses at the IS and terminal had the same maximum amplitude (4.45 mV) and t_1/2_ value (9.07 ms). These rise times are shorter than those observed experimentally, likely because the latter incorporates voltage-gated channel kinetics (Barnes, 1994; Van Hook et al., 2019).

**Figure 3.**
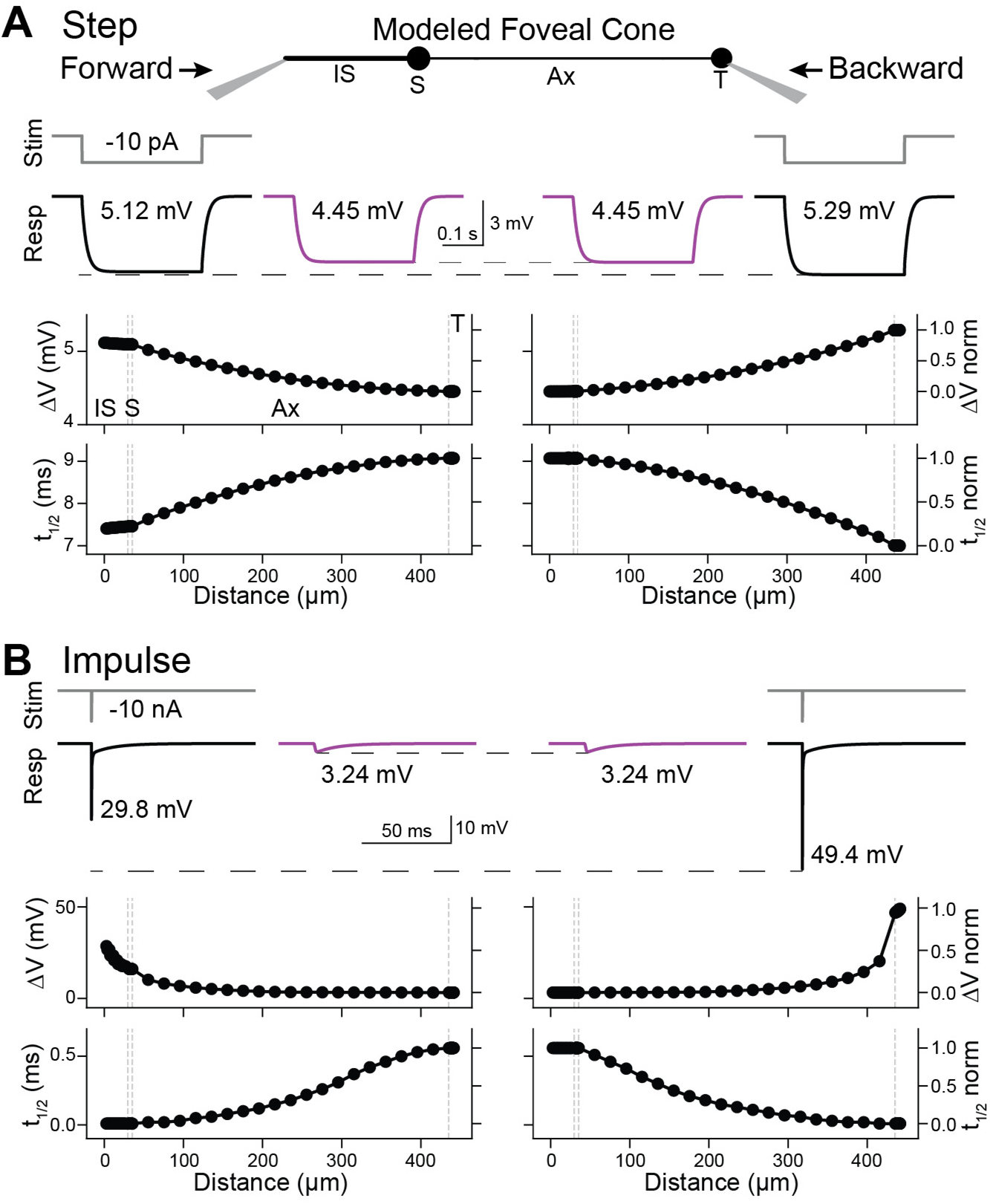
Backpropagation of responses to steps in a passive model of a foveal cone. **A.** Current steps (10 pA, 300 ms) were injected into the inner segment (left) or terminal (right) of a passive, compartmental model of a foveal cone (center, with compartments labeled: IS, inner segment; S, soma; Ax, axon; and T, terminal). Local (black, with stimulus monitor in gray) and propagated (magenta) voltage responses are shown for the most distal tips of the IS and T, and the maxima noted. The maximum voltage change and time to half-maximum (t_1/2_) are plotted across the length of the model cell. Gray dashed lines mark the borders of cell compartments. **B.** As for the plots in **A**, but for responses to impulse stimuli (10 nA, 0.01 ms). See **Figure S3B** for analyses in the frequency domain.

The reference foveal cone resembles actual cones in having a higher input resistance at the terminal than the IS (0.53 and 0.51 GΩ, respectively) and a larger local response at the terminal. The propagated/local response amplitude ratios were 0.84 (backward) and 0.87 (forward), and the local/propagated t_1/2_ ratios were 0.77 (backward) and 0.82 (forward). These amplitude and kinetic ratios are smaller than those we observed experimentally. The reason may be that the modeled cone represents a morphological extreme, while the recorded cones were varied (e.g., IS to terminal of 443 µm for the model cone and 205 ± 71 µm for the 10 recorded foveal cones). Indeed, shortening the model cone to 205 µm (axon length of 163 µm) resulted in higher ratios for response amplitude (0.96 and 0.97) and t_1/2_ (0.94 and 0.96) for backpropagation and forward propagation, respectively.

Impulse stimuli are often used to query the electrotonic properties of cells (Major et al., 1994). To situate our work in this context, we injected an essentially instantaneous current into the reference foveal cone (10 nA in 0.01 ms, the simulation time step; **Figure 3B**). Propagated impulse responses had identical peak amplitudes and t_1/2_ in the IS and terminal (3.24 mV and 0.56 ms), and local responses were larger for the terminal (49.4 mV) than the IS (29.8 mV, both with a 0.01-ms t_1/2_). In these respects, impulse and step responses are similar. They differ in that impulse responses, containing higher temporal frequencies, are attenuated more steeply during propagation (**Figure S3**).

Simulated responses to white noise (**Figure 4**) resembled empirical responses (**Figure 2**). One exception is that simulated phase delays were shorter; this is consistent with the briefer rise times for the step stimulus (see above) and the lack of voltage-gated channel activity in the model. This lack is also evident in the simpler shapes of the magnitude and phase spectra (**Figure S3**). Local responses were slightly larger at the terminal. For example, at 1 Hz, they were 0.53 and 0.51 GΩ. Accordingly, the propagated/local amplitude ratio appeared lower for backpropagation (0.84, 1 Hz) than forward propagation (0.87, 1 Hz). Nevertheless, across temporal frequencies, propagated responses had the same magnitude and phase in the terminal and IS (1, 60, and 100 Hz). The waveforms and features of linear filters calculated from these white noise simulations matched those of impulse responses (**Figure 3B** and **S3B, C**), as expected from a linear system.

**Figure 4.**
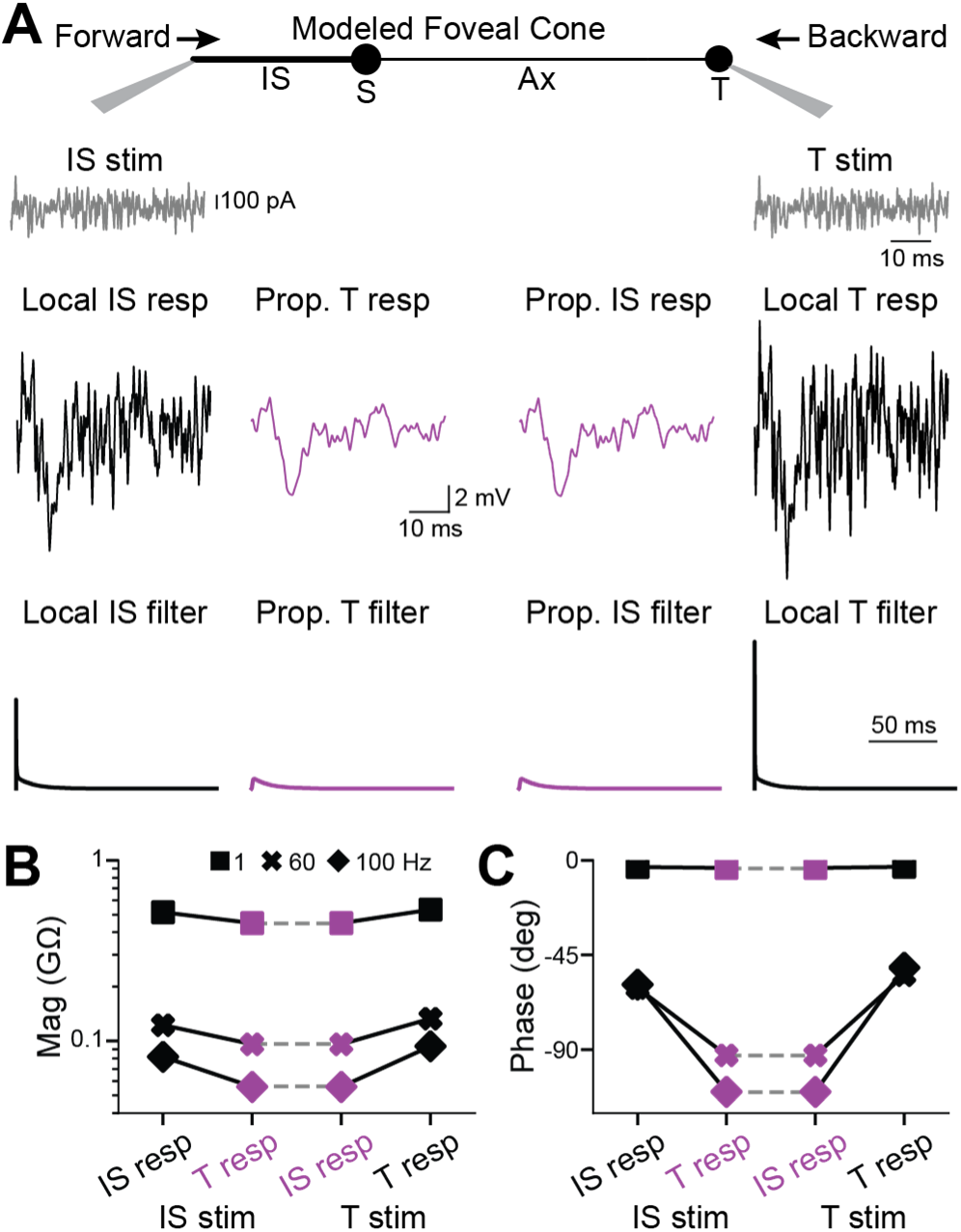
Backpropagation of responses to white noise in a passive model of a foveal cone. **A.** Current in the form of white noise was injected into the inner segment (left) or terminal (right) of a passive, compartmental model of a foveal cone. Local (black, with stimulus monitor in gray) and propagated (magenta) voltage responses are shown. Their linear filters are below. **B.** Response magnitudes at the inner segment and terminal, for forward and backward propagation, are shown for different temporal frequencies. See **Figure S3C** for further analyses of propagation across frequencies. **C.** As in **B** but for response phase.

To summarize, our simulations indicate that passive response propagation through foveal cones is effective in both forward and backward directions, whether the responses were evoked by step, impulse, or white-noise stimuli.

### Backpropagation to the phototransducing outer segment

The phototransducing outer segment (OS) may influence response propagation. This compartment is lost from our dissociated cones, so we studied it using the reference foveal cone model. In prior studies, which concerned forward propagation, the OS was modeled minimally as a resistor, capacitor, and battery (Hsu et al., 1998; Bryman et al., 2020). Here, we modeled a compartment of length and diameter reflecting measured morphology (Dowling, 1965) (**Materials and Methods**). We also considered the membrane discs within the OS, some of which appear to be continuous with the plasma membrane and therefore add to its capacitance and conductivity (Cohen, 1961; Dowling, 1965; Anderson and Fisher, 1979). An early, electron-microscopic study suggested that 10% of the discs are continuous in foveal cones (Cohen, 1961). However, this may be an underestimate because points of continuity could lie outside the single planes examined. We therefore assessed 10% and a likely overestimate of 60%. The connection between disc and plasma membrane has unknown geometry. Thus, we modeled the added surface area from continuous discs simply as increases in OS length or diameter.

Adding an OS lowered input resistance, response amplitudes, and t_1/2_ values. These reductions were higher with larger fractions of continuous discs. Nevertheless, whether delivering current in the form of a step, an impulse, or white noise, both forward (not shown) and backward (**Figure 5A**) propagation remained effective. This was the case whether we added disc membrane by lengthening or broadening the OS (**Figure 5B**). The influence of the OS on propagation effectiveness appears mild.

**Figure 5.**
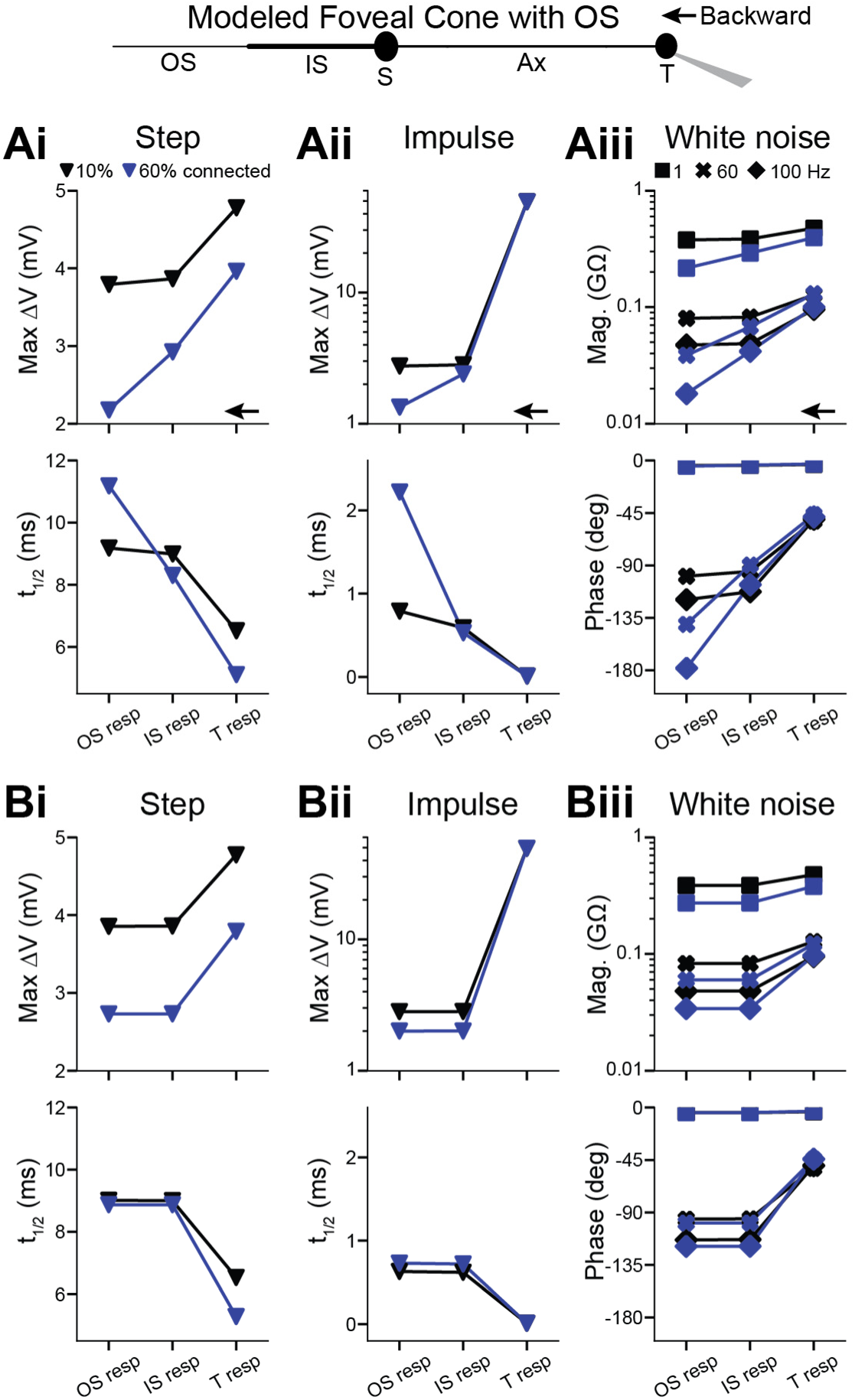
Backpropagation to the outer segment in a passive model of a foveal cone Ai. An outer segment (OS) was added to the passive, compartmental model of a foveal cone. The fraction of discs continuous with the plasma membrane was estimated to be 10% (black) or 60% (blue). The OS diameter was constant, and its length increased to account for the surface area of attached discs. The terminal received a current step (10 pA, 300 ms) and responses given at the terminal, IS, and OS. **Aii.** As in **Ai** but for an impulse stimulus (10 nA, 0.01 ms). **Aiii.** As in **Ai** but for white noise. **Bi-iii.** As in **Ai-iii** but keeping OS length constant and increasing diameter to account for the surface area of continuous discs.

### Backpropagating signals from network inputs

The high fidelity of backpropagation raises the possibility that inputs to the terminals of foveal cones may influence the OS voltage and thus the ionic currents of phototransduction. To investigate this possibility, we first considered gap-junctional coupling of foveal cone terminals (Raviola and Gilula, 1973; Ahnelt and Pflug, 1986; Tsukamoto et al., 1992; DeVries et al., 2002; Kim et al., 2025). We simulated an array of reference foveal cones, which comprised a central cone and 3 concentric rings of surrounding cones arranged in a naturalistic, hexagonal pattern (Schein, 1988). We tested gap-junctional conductances spanning a likely range (0.1 and 1 nS)(DeVries et al., 2002; Hornstein et al., 2004; Sinha et al., 2017).

We asked how stimulation of a single cone influences a neighboring cone (**Figure 6A**). We delivered current to the OS of the “sending” cone. We measured the propagated response at the “receiving” cone’s IS/OS junction and OS tip (length increased to mimic 10% continuous discs). The path of propagation is therefore forward through the sending cone and backward through the receiving cone, with the local response at the sending IS and the propagated responses at the receiving IS/OS junction and distal OS. Forward and backward propagation through coupled cones was effective for step, impulse, and white noise stimuli. The higher gap-junctional conductance produced a lower input resistance and thus responses that were smaller and faster. Nevertheless, propagation effectiveness was unaffected.

**Figure 6.**
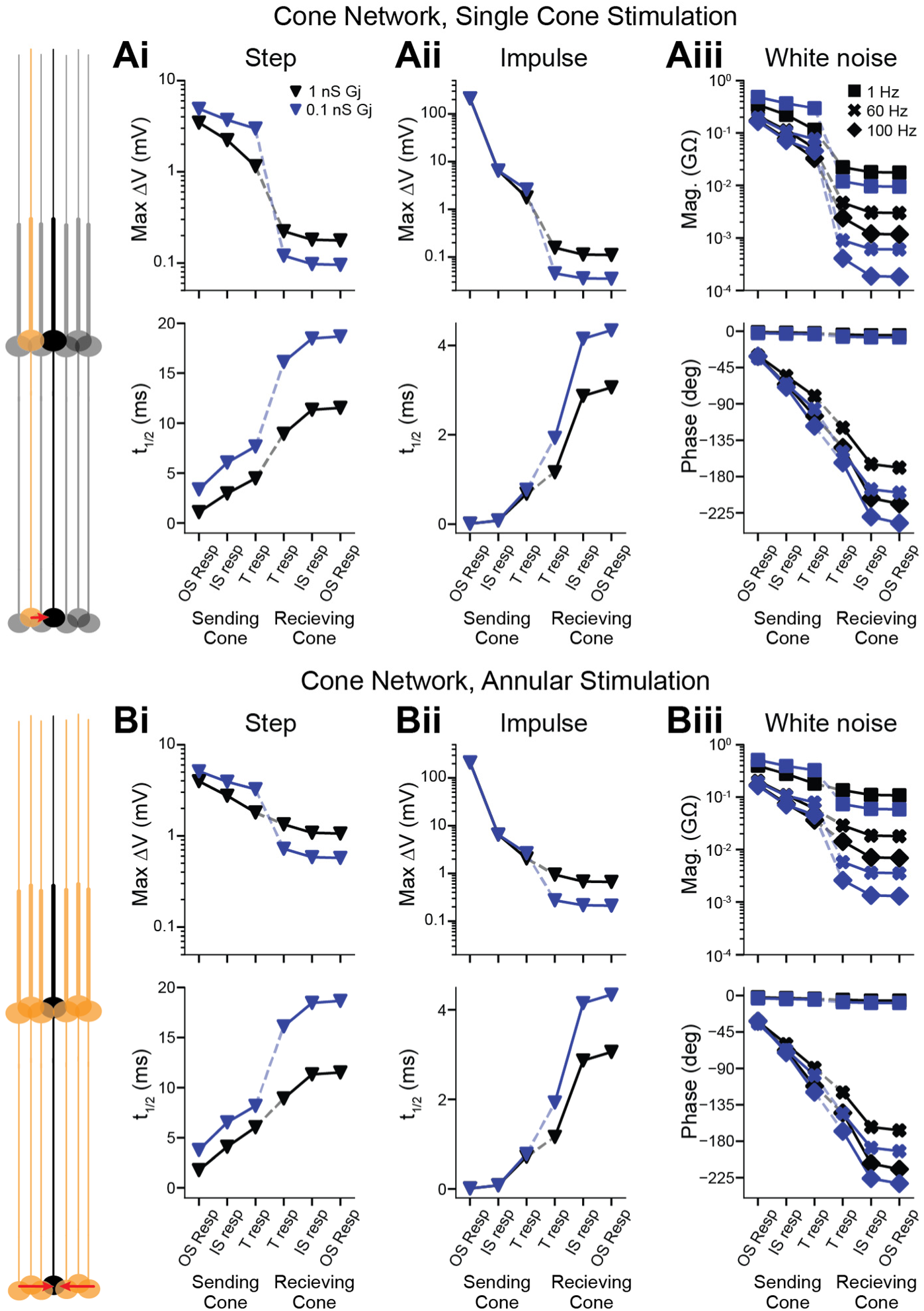
Backpropagation of gap-junctional inputs to the outer segment in cone networks. **A.** The model comprised a central cone and three concentric rings of additional cones in a hexagonal array. Neighboring cones are connected via gap junctions. A stimulus was given to the OS of one cone (“sending” cone), and the response followed through subsequent compartments to the OS of the central cone (“receiving” cone). The dashed line indicates the gap junction. Shown are simulated responses to steps (10 pA, 300 ms; **Ai**), impulses (10 nA, 0.01 ms, **Aii**), and white noise (**Aiii**). Gap-junction conductances were 1 nS (black) or 0.1 nS (blue). 10% of OS discs are continuous with the plasma membrane (modeled by OS lengthening). **B.** As in **A** but for 6 sending cones in an annulus around the receiving cone. Responses are identical across sending cones (one shown).

Most response attenuation and slowing took place at the gap junctions. For example, given the step stimulus (10 pA, 300 ms) and a coupling conductance of 1 nS, the sending cone’s distal OS, IS/OS junction, and terminal had responses of 3.45, 2.22 and 1.15 mV, and the receiving cone’s terminal, IS/OS junction, and distal OS had responses of 0.222, 0.180, and 0.177 mV. The maximum response of a cone is ∼4-fold larger and would therefore produce propagated responses of ∼0.7 mV at the IS/OS junction and distal OS.

A larger propagated response would be expected if the receiving cone were surrounded by sending cones. We produced a model in which the receiving cone was the center of a ring of sending cones, which was itself surrounded by two rings of unstimulated cones (**Figure 6B**). Adding even more rings had negligible effects, as responses were transmitted largely between adjacent cones (∼21% transmitted from the sending cone its neighbor) and scarcely beyond (∼8% from the sending cone to a second-order neighbor) at the highest tested coupling conductance. The propagated responses in the receiving cone were 6-fold higher with annular stimulation than single-neighbor stimulation, consistent with linear summation in the hexagonal cone array. Considering a saturated response in each sending cone (40 pA, which generates a ∼20 mV response locally), the 1-nS coupling conductance gives responses in the receiving cone of 4.32 mV at the IS/OS junction and 4.24 mV at the distal OS (see below for the likely influence of this response). Response latencies for annular stimulation were identical to those of single-cone stimulation (Luo et al., 2008).

To investigate additional network inputs, we turned to published recordings of macaque cones stimulated at their center (a spot of light) or surround (an annulus of light)(Verweij et al., 2003). The former produced a net outward current, attributed to phototransduction. The latter produced a net inward current, attributed to horizontal-cell feedback, which was larger for annuli of greater outer diameters. We simulated the center current at the OS, either alone or in combination with a surround current at the terminal. At the largest annulus size, we found that the sum gave a maximum depolarization of 8.59 mV at the terminal and, following backpropagation, 6.71 mV at the OS (**Figure 7**; we provide an interpretation of this response below).

**Figure 7.**
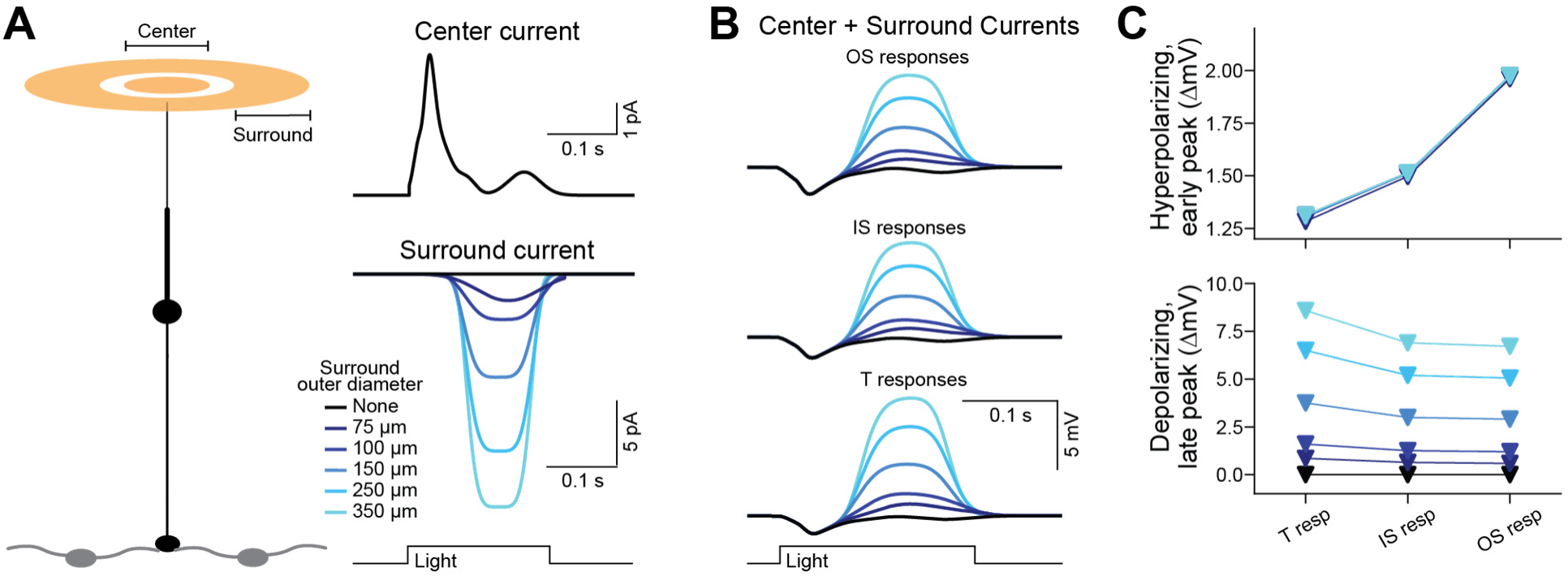
Propagation of simulated center and surround light responses. **A.** *Left,* schematic of the previously published experiment using center and surround illumination (Verweij et al., 2003). *Right,* Simulations of currents evoked by center (top) or surround (bottom) illuminations. The center stimulus was a 35-µm diameter spot. The surround stimulus was an annulus with a 40-µm inner diameter and varied outer diameters (75-350 µm). **B.** The center current was given at the distal OS and each surround current at the terminal of the reference foveal cone. Voltage responses are shown at the distal OS, OS/IS junction, and terminal. 10% of the OS discs are assumed to be continuous with the plasma membrane (modeled by OS lengthening). **C.** Center and surround response amplitudes as a function of subcellular location.

## DISCUSSION

We have investigated how signals backpropagate through cone photoreceptors of the primate fovea, from the presynaptic terminal to the outer segment (OS), and how such signals may shape the encoding of visual information.

### Passive backpropagation

We found that backpropagation is highly effective in foveal cones despite their length, slenderness, and modestly-sized presynaptic terminal. Active properties appear unnecessary. Indeed, our passive model shows backpropagation that is on par with that of actual foveal cones. This model has biophysical properties, measured from foveal cones, that are suited to propagation. The first is a low intracellular resistivity that facilitates current flow within the cell (Bryman et al., 2020). Intracellular resistivity has been examined electrophysiologically in turtle cones and appears high (Lasater et al., 1989). Nevertheless, the signal amplitude drops only 3% during backpropagation, likely because the axon is short. Thus, cones may achieve effective backpropagation through different combinations of biophysical and morphological properties.

Foveal cones also have a high membrane resistivity, which promotes propagation by limiting current loss from the cell (Bryman et al., 2020). This property distinguishes foveal from peripheral cones and is largely explained by smaller voltage-gated currents in the former, mostly a hyperpolarization-activated cation current (I_H_) and a K^+^ current (I_Kx_) (Bryman et al., 2020). These currents, which are active across the cone’s physiological range of membrane voltage and do not inactivate, contribute to the cell’s passive properties. Boosting propagation by reducing voltage-gated current is opposite to the common case of increasing them for amplification (Hille, 2001; Bean, 2007; Carter et al., 2012).

Some neurons are analogous to foveal cones in favoring backpropagation via a reduction of voltage-gated current. For example, pyramidal neurons express a dendritic K^+^ current, I_A_, which is reduced by neuromodulation and long-term plasticity to increase backpropagation (Hoffman et al., 1997; Hoffman and Johnston, 1999; Kim et al., 2007). Future studies might investigate the possibility of such modulation in primary sensory neurons like foveal cones.

Backpropagation through foveal cones may not require voltage-gated currents but we do observe signs of their activity, such as bandpass tuning to temporal frequency. Nevertheless, the backpropagating response resembles that of a passive system in being accurately described as a linear filter. This linearity facilitates prediction of foveal cone responses. The role of voltage-gated currents in foveal cones remains an open question; in cones of the peripheral retina and other species, these currents are involved in shaping response kinetics, improving signal to noise, and mediating adaptation (Barnes, 1994; Pahlberg et al., 2017; Van Hook et al., 2019; Saha et al., 2024).

Much remains to be learned about the identities of ion channels in foveal cones. By analogy to cones and rods examined across retinal regions and species, they are likely to represent major types that are gated by voltage directly as well as indirectly, such as in the case of Ca^2+^-activated Cl^-^ channels (Yagi and Macleish, 1994; MacLeish and Nurse, 2007; Frederiksen et al., 2025). Channels that are unorthodox in photoreceptors, such as voltage-gated Na^+^ channels, might also be considered (Ohkuma et al., 2007). Answers are nearer at hand given the introduction of single-cell transcriptomics to the fovea (Peng et al., 2019; Yan et al., 2020) and continued refinements in protein characterization within primate retina (Gayet-Primo et al., 2018).

### Backpropagation in the context of phototransduction

Effective backpropagation raises the possibility that inputs to the terminals of foveal cones could substantially alter the driving force for transmembrane Ca^2+^ flux at the distal OS, thus setting the gain and kinetics of phototransduction (Yau, 1991; Burns and Baylor, 2001). Our simulations suggest that this may not be the case, given the estimated size of terminal inputs and the voltage-dependence of Ca^2+^ flux through the phototransduction cascade’s CNG channels (Ohyama et al., 2002). We estimate that gap-junction inputs could hyperpolarize the OS by as much as ∼4 mV, while horizontal cell feedback could depolarize the OS by ∼7 mV. While these voltage changes are substantial given that the light response itself is only tens of millivolts, they are likely to change Ca^2+^ flux through the transduction channels by only 0.04 and 0.03 pA, respectively (Ohyama et al., 2002). Thus, even though backpropagated responses are scarcely attenuated, information flow through foveal cones may remain unidirectional.

Our study has several caveats. First, we examined gap junctions among terminals of neighboring foveal cones, and the junctional conductances have not been defined. We tested a range that is likely physiological (Hornstein et al., 2004). Actual conductances may be the smaller side (Angueyra and Rieke, 2013; Sinha et al., 2017). Hence, our estimates of gap junctional influences are likely to be upper bounds. A second caveat is that, to investigate surround inhibition, we used currents measured in peripheral cones (Verweij et al., 2003). They have not been measured for foveal cones. This inhibition is likely to be larger in the periphery due to the greater signal pooling there (Boycott and Kolb, 1973). Therefore, the effect we observe is likely to be an upper bound for foveal cones. A third caveat is that we have not examined other potential inputs to foveal cone terminals, such as current from electrogenic glutamate transport (Thoreson and Mangel, 2012; Hays et al., 2021). Fourth, the voltage-dependence of Ca^2+^ flux through native phototransduction channels has only been determined for teleost cones (Ohyama et al., 2002). If it is steeper for primate foveal cones, backpropagated responses from the terminal may indeed influence phototransduction. As these parameters come to be defined, our single-cell and network models provide frameworks for examining their influences.

Foveal cones have slower light responses than peripheral cones (Sinha et al., 2017; Baudin et al., 2019; Bryman et al., 2020; Saha et al., 2024). An explanation was that slow responses are needed to avoid steep attenuation during forward propagation—such attenuation was likely given the elongated morphologies of these neurons as well as intracellular and membrane resistivities inferred from other cell types (Hsu et al., 1998; Masland, 2017). However, we now understand that the light response could be much faster and still propagate effectively from OS to terminal (Bryman et al., 2020). The reason for slow foveal phototransduction is therefore enigmatic. Backpropagation may provide a clue. Its fidelity is high overall but highest for lower frequencies. If phototransduction contained lower temporal frequencies than terminal inputs, foveal cones would be biased for the forward flow of information despite the bidirectional effectiveness of response propagation.

## AUTHOR CONTRIBUTIONS

S.R.W. analyzed the data, designed and performed the computational modeling, and interpreted the results. G.S.B. designed and conducted the experiments, collected tissue, built the original foveal cone model, and contributed to the analysis framework. M.T.H.D. conceived the study, designed the experiments, collected tissue, provided guidance on the modeling, and interpreted the results. S.R.W. and M.T.H.D. wrote the paper.

## ACKNOWLEDGMENTS

Andreas Liu and Michael Brown assisted with tissue collection. Stephen Massey, Samuel Brill-Weil, Franklin Caval-Holme, Philippe Morquette, Navid Mousavi, and Viet Nguyen-Minh provided valuable discussion. Support was provided by the Helen Hay Whitney Foundation and National Heart, Lung, and Blood Institute grant T32HL007901 (S.R.W.); a National Science Foundation GRFP (G.S.B.); National Eye Institute grants EY025840, EY030628, EY025555, and EY028633 (M.T.H.D); the Harvard/MIT Joint Research Grants Program in Basic Neuroscience and BrightFocus Foundation grants (M.T.H.D.); National Eye Institute grant EY012196 (Harvard Medical School); and National Institute of Child Health and Human Development grant 1U54HD090255 (Boston Children’s Hospital).

## CONFLICT OF INTERESTS

M.T.H.D. is affiliated with the Center for Brain Science (Harvard University), the Division of Sleep Medicine (Brigham and Women’s Hospital, Harvard Medical School), and the Broad Institute of MIT and Harvard. All authors declare no competing interests.

---

**Figure S1.**
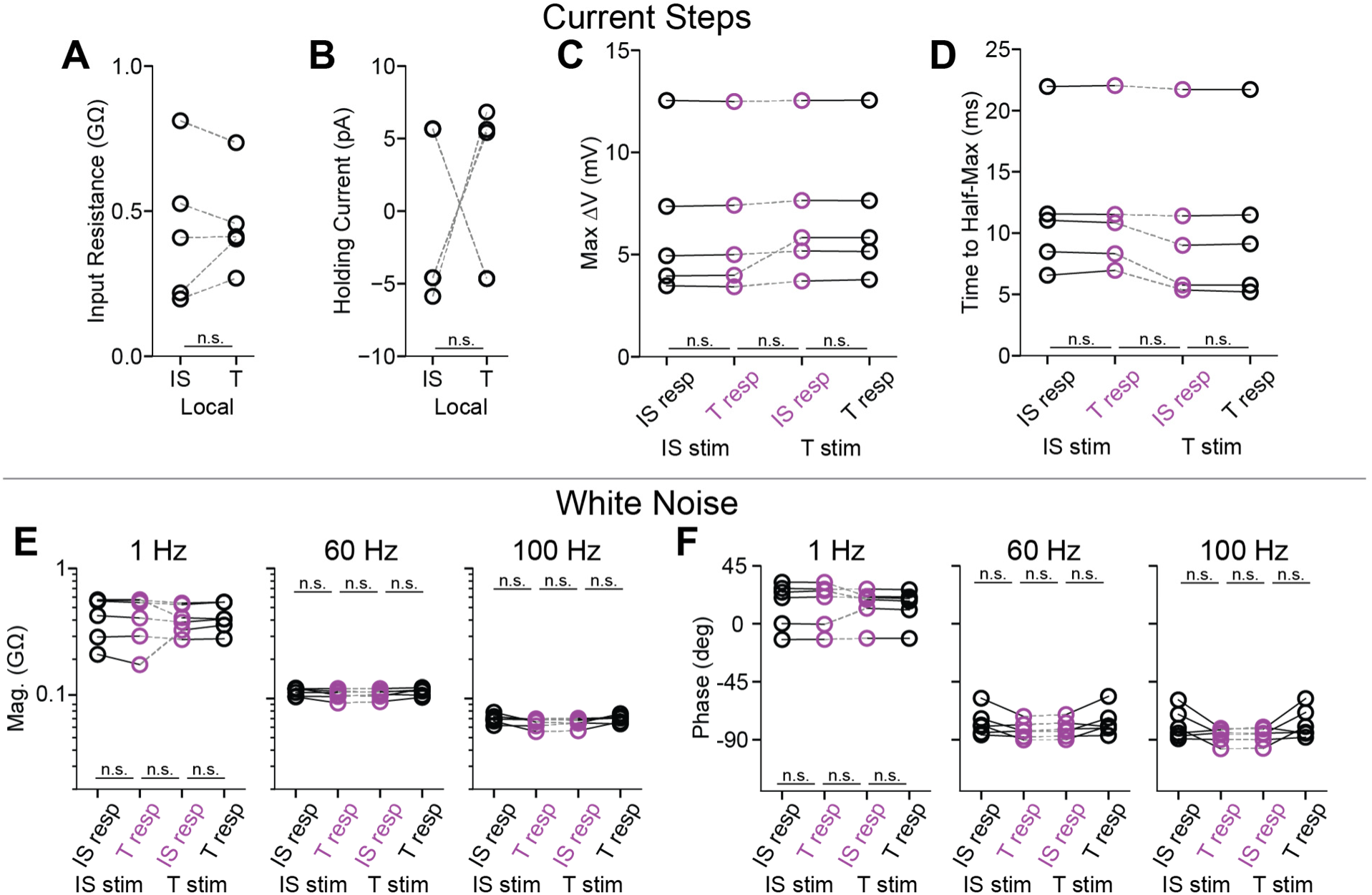
Propagation of steps and white noise in dissociated peripheral cones (related to Figures 1 and 2). **A.** Input resistances for the inner segment and terminal of each peripheral cone (calculated from local responses). **B.** Steady current injected into each compartment to maintain its membrane voltage near -60 mV. **C.** Maximum voltages of forward (IS stim) and backward (T stim) propagated responses to steps (-10 pA, 300 ms). Local (black) and propagated (magenta) response magnitudes are given (5 cells, Bonferroni correction for 3 comparisons). **D.** As in **C**, but for time to half-maximum. **E.** Magnitude of local (black) and propagated (magenta) responses to white noise (6 cells, Bonferroni correction for 3 comparisons). **F.** As in **E**, but for phase.

**Figure S2.**
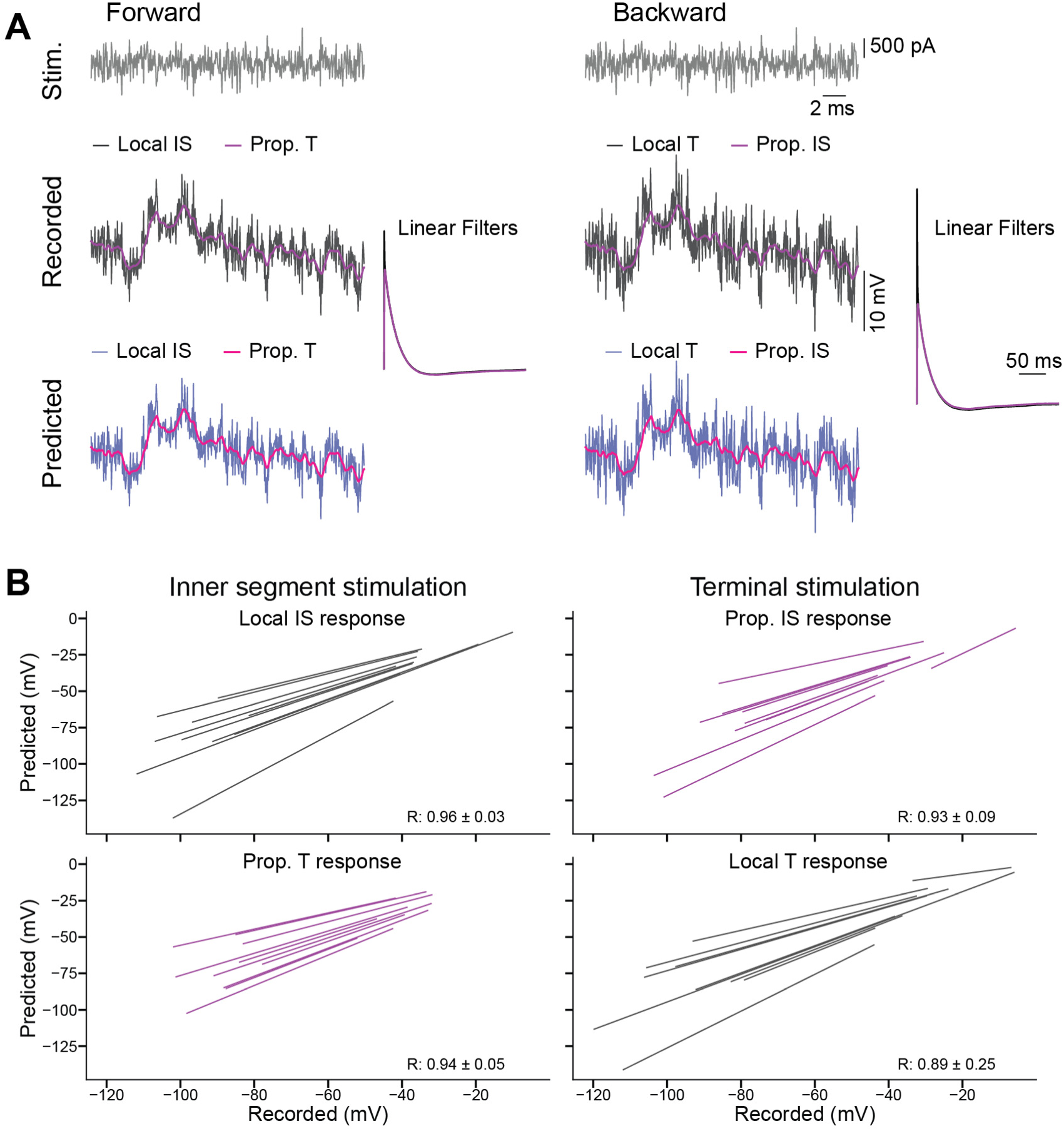
Filter verification and response linearity in foveal cones (related to Figure 2). **A.** White noise current was given (gray) to either the IS (*left*, forward propagation) or terminal (*right*, backward propagation) and voltage responses measured (middle). Linear filters were calculated (right) and convolved with the stimulus to generate predicted responses (bottom). This was done for forward (left) and backward (right) propagation and for local (black) and propagated (magenta) responses. Filters were calculated from one response segment and tested on held-out segments. **B.** Comparing the relationship between recorded and predicted responses for forward (left) and backward (right) propagation (10 cells). The average Pearson’s R and standard deviation are reported for each condition (R=0.923 ± 0.093, grand mean ± SD).

**Figure S3.**
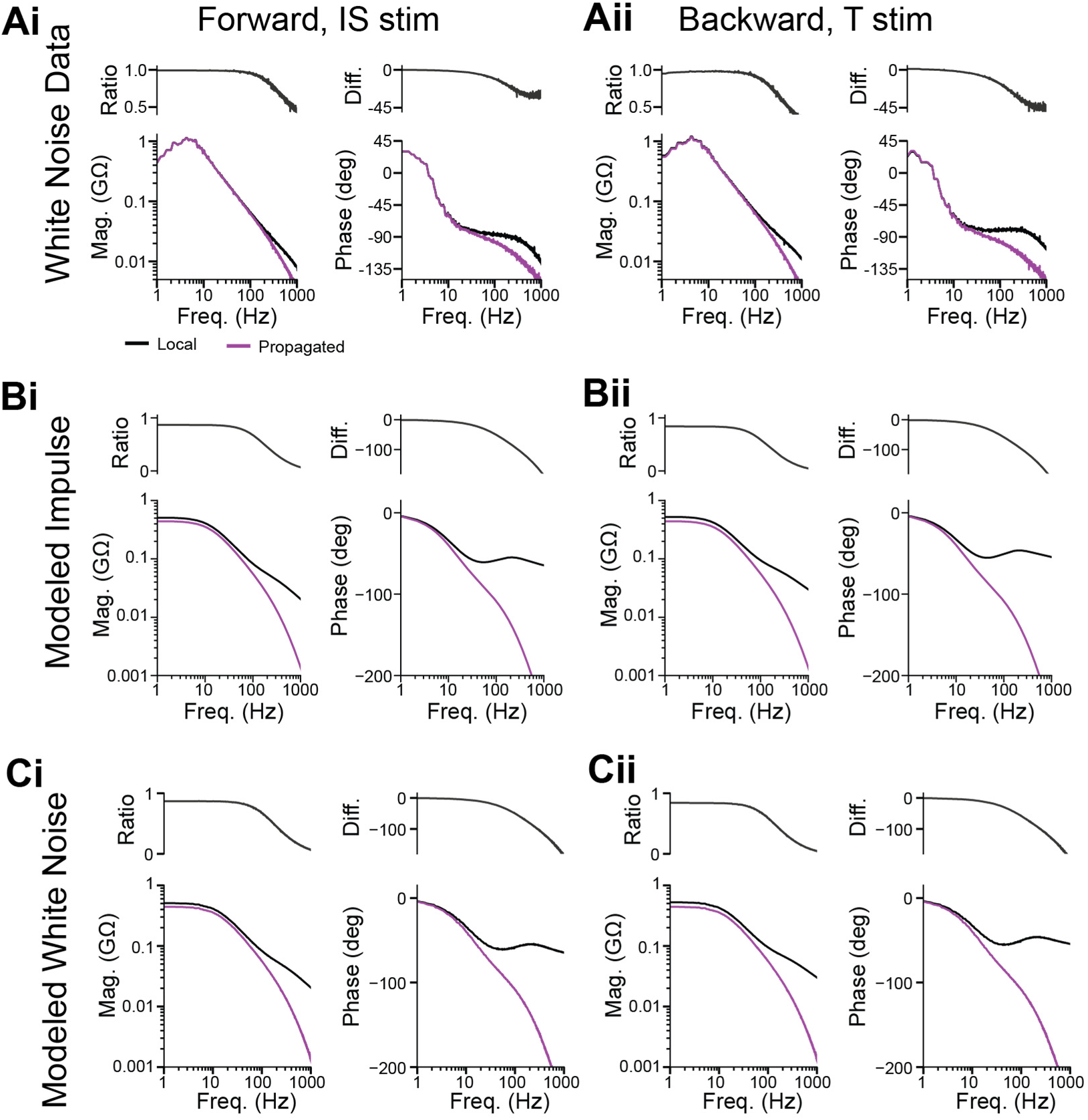
Comparison of measured and modeled foveal cone responses over an extended frequency range (related to Figures 2-4). **A.** White noise was delivered to a dissociated foveal cone at the IS (i) or terminal (ii) for evaluation of forward and backward propagation, respectively. Magnitude and phase spectra are shown for local (black) and propagated (magenta) responses over an extended range of temporal frequencies. Spectra are smoothed using a moving median filter (3-sample window). Magnitude ratios and phase differences, comparing local and propagated responses, are shown. Same example cell as the cell shown in Fig. 2B. **B.** As in **A** but for the reference foveal cone model given impulse stimuli. **C.** As in **A** but for the reference foveal cone model given white noise stimuli.

**Table S1.** Parameters of responses to steps in foveal and peripheral cones (related to Figure 1).

|  | Foveal cones, N=9 cells |  |  |  |  |  |  |  |
| --- | --- | --- | --- | --- | --- | --- | --- | --- |
|  | IS stim. - Forward |  |  |  | Terminal stim. - Backward |  |  |  |
|  | IS response |  | Terminal response |  | IS response |  | Terminal response |  |
| Parameter | Mean | SD | Mean | SD | Mean | SD | Mean | SD |
| Baseline (mV) | -59.57 | 6.17 | -59.41 | 6.85 | -59.52 | 6.00 | -59.36 | 6.62 |
| Peak response (mV) | -7.67 | 1.42 | -7.44 | 1.46 | -7.49 | 1.46 | -8.04 | 1.36 |
| Steady-state response (mV) | -5.02 | 1.42 | -4.81 | 1.58 | -4.95 | 1.58 | -5.50 | 1.53 |
| Time to 5% peak (ms) | 1.03 | 0.55 | 1.83 | 0.73 | 1.64 | 0.45 | 0.74 | 0.37 |
| Time to 50% peak (ms) | 13.88 | 2.38 | 14.83 | 2.18 | 14.56 | 1.91 | 13.57 | 2.12 |
| Time to peak (ms) | 70.58 | 14.64 | 71.32 | 15.88 | 72.67 | 11.35 | 72.75 | 11.31 |
| Input resistance (GΩ) | 0.50 | 0.14 |  |  |  |  | 0.55 | 0.15 |
| Holding current (pA) | -0.32 | 5.70 |  |  |  |  | 1.34 | 5.66 |
|  | Peripheral cones, N=5 cells |  |  |  |  |  |  |  |
|  | IS stim. - Forward |  |  |  | Terminal stim. - Backward |  |  |  |
|  | IS response |  | Terminal response |  | IS response |  | Terminal response |  |
| Parameter | Mean | SD | Mean | SD | Mean | SD | Mean | SD |
| Baseline (mV) | -60.61 | 5.09 | -59.94 | 6.19 | -60.56 | 5.23 | -59.89 | 6.27 |
| Peak response (mV) | -6.45 | 3.72 | -6.46 | 3.70 | -6.98 | 3.42 | -6.99 | 3.41 |
| Steady-state response (mV) | -3.78 | 2.55 | -3.79 | 2.53 | -4.52 | 1.71 | -4.54 | 1.71 |
| Time to 5% peak (ms) | 0.74 | 0.62 | 0.77 | 0.67 | 0.70 | 0.58 | 0.71 | 0.63 |
| Time to 50% peak (ms) | 11.92 | 5.96 | 11.94 | 5.94 | 10.65 | 6.67 | 10.66 | 6.69 |
| Time to peak (ms) | 59.27 | 23.10 | 59.46 | 23.43 | 117.97 | 77.70 | 118.22 | 77.60 |
| Input resistance (GΩ) | 0.43 | 0.25 |  |  |  |  | 0.46 | 0.15 |
| Holding current (pA) | -0.75 | 5.87 |  |  |  |  | 1.71 | 5.83 |

**Table S2.** Statistics for responses to steps in foveal and peripheral cones (related to Figure 1). *. Comparisons were made for local responses only.

|  | Foveal cones, N=9 cells |  |  |  |  |  |
| --- | --- | --- | --- | --- | --- | --- |
|  | IS stim., IS vs. Terminal resp. |  | Terminal stim., IS vs. Terminal resp. |  | Propagated responses |  |
| Parameter | p value | Cohen's $\delta$ | p value | Cohen's $\delta$ | p value | Cohen's $\delta$ |
| Baseline (mV) | 1.0000 | 0.0244 | 1.0000 | 0.0256 | 1.0000 | 0.0169 |
| Peak response (mV) | 0.0586 | 0.1631 | 0.0234 | 0.3900 | 1.0000 | 0.0312 |
| Steady-state response (mV) | 0.0586 | 0.1429 | 0.0234 | 0.3538 | 1.0000 | 0.0916 |
| Time to 5% peak (ms) | 0.0234 | 1.2421 | 0.0234 | 2.1698 | 1.0000 | 0.3153 |
| Time to 50% peak (ms) | 0.0234 | 0.4169 | 0.0234 | 0.4920 | 1.0000 | 0.1279 |
| Time to peak (ms) | 1.0000 | 0.0483 | 1.0000 | 0.0071 | 1.0000 | 0.0982 |
| Input resistance (GΩ)* | 0.0391 | 0.3289 | | | | |
| Holding current (pA)* | 0.8203 | 0.2908 |  |  |  |  |
|  | Peripheral cones, N=5 cells |  |  |  |  |  |
|  | IS stim., IS vs. Terminal resp. |  | Terminal stim., IS vs. Terminal resp. |  | Propagated responses |  |
| Parameter | p value | Cohen's $\delta$ | p value | Cohen's $\delta$ | p value | Cohen's $\delta$ |
| Baseline (mV) | 1.0000 | 0.1172 | 1.0000 | 0.1156 | 1.0000 | 0.1067 |
| Peak response (mV) | 1.0000 | 0.0025 | 1.0000 | 0.0027 | 0.1875 | 0.1454 |
| Steady-state response (mV) | 1.0000 | 0.0026 | 0.9375 | 0.0080 | 0.9375 | 0.3413 |
| Time to 5% peak (ms) | 0.8551 | 0.0495 | 1.0000 | 0.0265 | 1.0000 | 0.1149 |
| Time to 50% peak (ms) | 1.0000 | 0.0027 | 1.0000 | 0.0012 | 0.1875 | 0.2040 |
| Time to peak (ms) | 1.0000 | 0.0079 | 0.3750 | 0.0033 | 0.9375 | 1.0196 |
| Input resistance (GΩ)* | 0.8125 | 0.1083 | | | | |
| Holding current (pA)* | 1.0000 | 0.4203 |  |  |  |  |

**Table S3.** Parameters of responses to white noise in foveal and peripheral cones (related to Figures 2 and S1).

|  | Foveal cones, N=10 |  |  |  |  |  |  |  |
| --- | --- | --- | --- | --- | --- | --- | --- | --- |
|  | IS stim. |  |  |  | Terminal stim. |  |  |  |
| Quantifying | IS response |  | Terminal response |  | IS response |  | Terminal response |  |
| Magnitude (GΩ) | Mean | SD | Mean | SD | Mean | SD | Mean | SD |
| 1 Hz | 0.419 | 0.152 | 0.405 | 0.154 | 0.422 | 0.189 | 0.453 | 0.179 |
| 60 Hz | 0.096 | 0.011 | 0.090 | 0.013 | 0.090 | 0.014 | 0.101 | 0.010 |
| 100 Hz | 0.062 | 0.008 | 0.054 | 0.009 | 0.054 | 0.009 | 0.067 | 0.008 |
| Phase (deg) |  |  |  |  |  |  |  |  |
| 1 Hz | 20.21 | 9.53 | 21.46 | 11.06 | 19.02 | 10.48 | 16.70 | 8.65 |
| 60 Hz | -78.85 | 8.64 | -91.82 | 6.42 | -91.71 | 6.97 | -74.99 | 10.68 |
| 100 Hz | -80.87 | 11.06 | -100.14 | 11.00 | -99.35 | 10.46 | -75.02 | 13.30 |
|  | Peripheral cones, N=6 |  |  |  |  |  |  |  |
|  | IS stim. |  |  |  | Terminal stim |  |  |  |
| Quantifying | IS response |  | Terminal response |  | IS response |  | Terminal response |  |
| Magnitude (GΩ) | Mean | SD | Mean | SD | Mean | SD | Mean | SD |
| 1 Hz | 0.441 | 0.154 | 0.431 | 0.162 | 0.414 | 0.103 | 0.428 | 0.104 |
| 60 Hz | 0.112 | 0.008 | 0.108 | 0.009 | 0.108 | 0.009 | 0.113 | 0.007 |
| 100 Hz | 0.070 | 0.006 | 0.065 | 0.006 | 0.065 | 0.005 | 0.070 | 0.005 |
| Phase (deg) |  |  |  |  |  |  |  |  |
| 1 Hz | 15.41 | 17.53 | 15.52 | 17.66 | 14.71 | 13.62 | 14.00 | 13.44 |
| 60 Hz | -76.77 | 10.33 | -82.45 | 6.59 | -81.68 | 7.10 | -76.07 | 10.60 |
| 100 Hz | -78.62 | 11.68 | -86.87 | 5.85 | -86.61 | 6.04 | -77.36 | 11.61 |

**Table S4.** Statistics for responses to white noise (related to Figures 2 and S1).

|  | Foveal cones, N=10 cells |  |  |  |  |  |
| --- | --- | --- | --- | --- | --- | --- |
| Statistics | IS stim., IS vs. Terminal resp. |  | Terminal stim., IS vs. Terminal resp. |  | Propagated responses |  |
| Magnitude (GΩ) | p value | Cohen's δ | p value | Cohen's δ | p value | Cohen's δ |
| 1 Hz | 0.0586 | 0.0959 | 0.0176 | 0.1671 | 1.0000 | 0.0993 |
| 60 Hz | 0.0059 | 0.5333 | 0.0117 | 0.9576 | 1.0000 | 0.0040 |
| 100 Hz | 0.0059 | 0.9345 | 0.0059 | 1.4842 | 1.0000 | 0.0196 |
| Phase (deg) |  |  |  |  |  |  |
| 1 Hz | 0.8262 | 0.1206 | 0.0117 | 0.2417 | 0.3164 | 0.2263 |
| 60 Hz | 0.0293 | 1.7028 | 0.0176 | 1.8544 | 1.0000 | 0.0172 |
| 100 Hz | 0.0293 | 1.7451 | 0.0176 | 2.0332 | 0.4805 | 0.0715 |
|  | Peripheral cones, N=6 cells |  |  |  |  |  |
| Statistics | IS stim., IS vs. Terminal resp. |  | Terminal stim., IS vs. Terminal resp. |  | Propagated responses |  |
| Magnitude (GΩ) | p value | Cohen's δ | p value | Cohen's δ | p value | Cohen's δ |
| 1 Hz | 1.0000 | 0.0605 | 0.2813 | 0.1325 | 0.9375 | 0.1230 |
| 60 Hz | 0.6563 | 0.5582 | 0.2813 | 0.5618 | 1.0000 | 0.0918 |
| 100 Hz | 0.4688 | 0.8582 | 0.2813 | 0.9476 | 1.0000 | 0.0989 |
| Phase (deg) |  |  |  |  |  |  |
| 1 Hz | 1.0000 | 0.0064 | 0.0938 | 0.0521 | 1.0000 | 0.0514 |
| 60 Hz | 0.4688 | 0.6547 | 0.4688 | 0.6226 | 0.2813 | 0.1110 |
| 100 Hz | 0.9375 | 0.8933 | 0.4688 | 0.9989 | 1.0000 | 0.0436 |

## Notes

https://github.com/wienbar/ConeBackpropagationCode_2026

### Competing Interest Statement

The authors have declared no competing interest.

### Summary of Updates

This version of the manuscript contains new simulations and analyses, with associated figures and text. Furthermore, small additions and corrections to existing figures and text have been made, and references have been added.

